# Double emulsions as a high-throughput enrichment and isolation platform for slower-growing microbes

**DOI:** 10.1101/2022.10.23.513397

**Authors:** Alexandra L. McCully, McKenna Loop Yao, Kara K. Brower, Polly M. Fordyce, Alfred M. Spormann

## Abstract

Our understanding of *in situ* microbial physiology is primarily based on physiological characterization of fast-growing and readily-isolatable microbes. Microbial enrichments to obtain novel isolates with slower growth rates or physiologies adapted to low nutrient environments are plagued by intrinsic biases for fastest-growing species when using standard laboratory isolation protocols. New cultivation tools to minimize these biases and enrich for less well-studied taxa are needed. In this study, we developed a high-throughput bacterial enrichment platform based on single cell encapsulation and growth within double emulsions (GrowMiDE). We showed that GrowMiDE can cultivate many different microorganisms and enrich for novel taxa that are never observed in traditional batch enrichments. For example, preventing dominance of the enrichment by fast-growing microbes due to nutrient privatization within the double emulsion droplets allowed cultivation of novel *Negativicutes* and *Methanobacteria* from stool samples in rich media enrichment cultures. In competition experiments between growth rate and growth yield specialist strains, GrowMiDE enrichments prevented competition for shared nutrient pools and enriched for slower-growing but more efficient strains. Finally, we demonstrated the compatibility of GrowMiDE with commercial fluorescence-activated cell sorting (FACS) to obtain isolates from GrowMiDE enrichments. Together, GrowMiDE + DE-FACS is a promising new high-throughput enrichment platform that can be easily applied to diverse microbial enrichments or screens.

## Introduction

Microbial physiology is largely based on studies using microbes isolated in pure cultures. Techniques for isolating novel microbes for the last 140 years involve recreating a favorable environment and flux of nutrients in the laboratory that mimics natural conditions and then attempting to recover the grown novel species. As a result, current methods are inherently biased towards fast-growing microbes that can outcompete other species for shared nutrients, even species with minor differences in maximal growth rate (μ_Max_) (**Fig. S1-S3**)[1, 2]. The missing slower-growing species might have unknown physiological strategies to adapt to their natural environment and that likely play critical roles in community resilience [3]. For example, growth efficiency (growth yield) over growth rates has been suggested as a survival strategy in low nutrient flux or spatially-structured environments including biofilms[1, 2, 4] and the marine subsurface environment (~30% of Earth’s biomass)[5–8]. Therefore, there is a critical need to develop new cultivation tools that minimize the bias for fast microbial growth rates.

Traditional methods that limit competition for finite resources (i.e., by nutrient privatization) typically rely on spatial separation and include dilution-to-extinction or plating for colony forming units[9, 10]. However, these techniques depend on cell abundances, are low-throughput, and are not conducive to recovering microbes that grow poorly at air-liquid interfaces, especially anaerobes. Droplet microfluidics approaches have emerged as a powerful high-throughput technique to cultivate novel microorganisms in isolated bioreactors each with precise distributions of reagents necessary for growth[10–14], facilitating screens for antibiotic resistances[11, 15, 16], and measurements of cellular activity[17–19]. Most microbiological droplet approaches use single emulsion droplets, in which aqueous droplets containing cells and medium are suspended within an oil phase. Many studies have shown the utility of single emulsion droplet microfluidics for surveying microbial diversity[19–22], obtaining high quality genomes[23], and functional screens[10, 11, 14, 16]. Single emulsion enrichment techniques have also demonstrated selection for strains that have elevated growth yields or slower growth rates, serving as a proof-of-concept for the impact of nutrient privatization on enrichment outcomes[24]. However, single emulsion droplets require custom equipment to analyze downstream and have only been sorted in proof-of-concept demonstrations with slow sorting speeds and limited fluorescence channels[25, 26], making them challenging to apply to new systems.

Double emulsion droplets (DEs) are an innovative microfluidics platform with the potential to greatly simplify microbial growth and isolations[27–30]. DEs consist of an inner aqueous compartment surrounded by an oil shell suspended within an outer aqueous layer (**Fig. 1B**). The outer aqueous suspension makes DEs directly compatible with common flow cytometry equipment, allowing them to be sorted via fluorescence-gated sorting (FACS) similar to cells[31]. Recent work optimizing device design, surfactants and sorting parameters has advanced the ability to sort DEs at high-throughput (12-14 kHz) and single droplet accuracy (>99% purity, 70% single droplet recovery) in traditional FACS instruments[31, 32]. Current applications for DE technology include cell encapsulation[19, 27], drug delivery[29], and protein function screens[19]. However, DEs have not yet been used as an enrichment platform for microorganisms.

**Figure 1.**
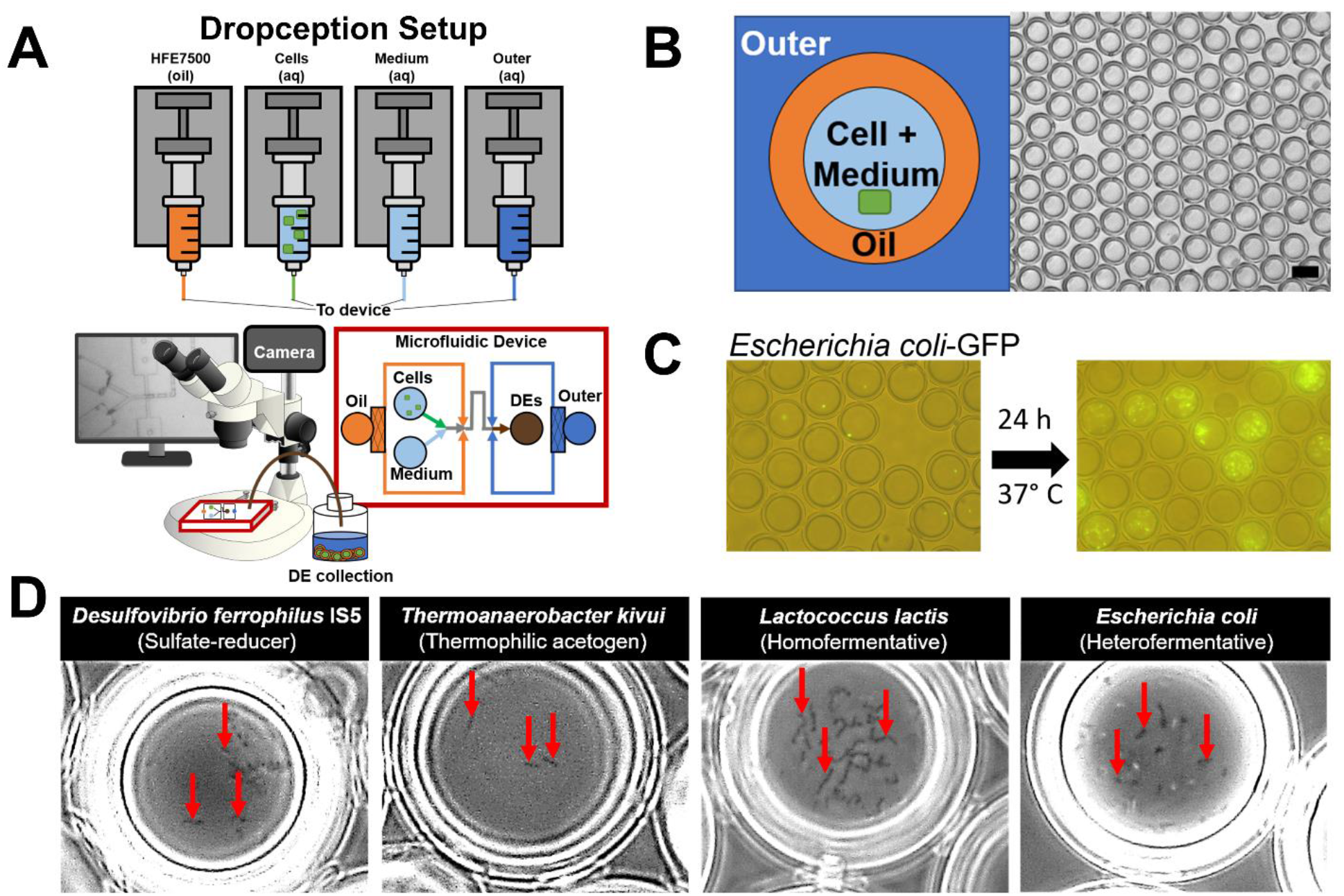
GrowMiDE: Double emulsion platform for bacterial cultivation. (**A**) Schematic representation of the GrowMiDE platform. Four syringe pumps drive oil and aqueous solutions into a custom microfluidic device to encapsulate single microbes within 30 or 45 μm diameter double emulsions (DEs); DE generation is monitored in real-time by a high-speed camera attached to an Amscope stereoscope. (**B**) Schematic and representative brightfield image of DE droplets. (**C**) Merged brightfield and fluorescent images of single *E. coli*-GFP cells loaded into DEs (left) and after growth in M9 + glucose for 24 hours (right). (**D**) Brightfield microscopy images indicating growth of diverse anaerobes within DEs including the sulfate-reducer *Desulfovibrio ferrophilus* IS5 on 60 mM lactate and 30 mM NaSO_4_, the acetogen *Thermoanaerobacter kivui* on 50 mM glucose at 65°C, lactic acid-producing fermenter *Lactococcus lactis* NZ9000 on 50 mM glucose, and mixed acid fermenter *E. coli* MG1655 on 50 mM glucose.

We designed a high-throughput DE platform (GrowMiDE) that i) is compatible with diverse microbial physiologies, ii) facilitates enrichments and subsequent isolations for downstream phenotypic characterizations, and iii) demonstrates recovery of microbial species that are normally lost due to outcompetition in batch culture enrichments. The GrowMiDE platform minimized bias for fast growth rates through nutrient privatization in individual droplets, facilitating enrichment of slower-growing species. Using GrowMiDE, we demonstrated enrichment of distinct community compositions compared to traditional batch cultures from the human gut microbiome, including a 22-fold increase in a novel Negativicutes species from the “most wanted” microbiome list[33]. We also determined that GrowMiDE can be applied to enrich for novel taxa due to prioritization of traits other than fast growth rates, specifically higher growth yields. Finally, we demonstrated the combination of GrowMiDE and DE-FACS as an isolation tool for microbiologists. This platform can be readily adapted to diverse biological systems and our results demonstrate the feasibility of high-throughput DE culturing to obtain cultured representatives of overlooked physiologies.

## Materials and Methods

### Strains and growth conditions

*E. coli* MG1655 and *E. coli*-GFP were routinely cultivated on LB agar or M9 medium supplemented with 25 mM glucose at 37°C with shaking. *Lactococcus lactis spp cremoris* WT (NZ9000) and the derived mutant ∆*ldhA* (NZ9010)[24] were cultivated in 10 mL of chemically defined medium (CDM)[34] supplemented with 1.5% casamino acids (w/v), 26 mg/L L-tryptophan, and 50 mM glucose at 30°C. Starter cultures of NZ9000 or NZ9010 were inoculated from a single colony from Difco M17 broth + 25 mM glucose (GM17) or GM17 + 5ug/mL erythromycin plates, respectively. *Pseudomonas putida* were routinely cultivated on LB agar or cetrimide plates. All growth curves were started with a 1% inoculum from a starter culture in stationary phase. For culturing strict anaerobes in DEs, 20 mL stationary cultures grown in anaerobic 60 mL serum vials were used as the inocula. *Desulfovibrio ferrophilus* IS5 (DSM no. 15579) was cultivated at 30°C in modified artificial seawater medium[35] supplemented with 60 mM lactate and 30 mM sulfate. *Thermoanaerobacter kivui* TKV002 was grown at 65°C in medium containing 25 mM MES (free acid), 75 mM MES (sodium salt), 13.7 mM NaCl, 0.8 mM MgSO_4_, 18.7 mM NH_4_Cl, 1.3 mM KCl, 0.1 mM CaCl_2_, 0.7 mM KH_2_PO_4_, 40 μM uracil, 1 mL/L trace element solution SL10, 1 mL/L selenate-tungstate solution, 1 mM Na_2_S, 0.5 mg/L resazurin, and 50 mM glucose as the catabolic substrate in DEs. Stool enrichments were grown at 37°C cultivated in modified mBHI medium containing: 37 g/L BHI mix (RPI), 200 mg/L tryptophan, 1 g/L arginine, 5 mg/mL menadione, 500 mg/L cysteine HCl, 0.12 μg/mL haemin solution[36], 0.2 g/L mucin, resazurin. For select enrichments, mBHI was supplemented with additional media components (mBHI^+^) based approximately on concentrations added to Gut Microbiota Medium (GMM)[36]: 3 mM sugars (glucose, cellobiose, maltose, fructose), 30 mM sodium acetate, 8 mM propionic acid, 4 mM sodium butyrate, 15 mM sodium lactate, 30 mM NaHCO_3_.

### Analytical procedures

Cell densities were determined based on the optical density at 600 nm (OD_600_) using an Ultraspec 2100 spectrophotometer (GE Healthcare) or a Tecan Infinite M1000 microplate reader. Glucose and fermentation profiles of *L. lactis* strains were quantified using an Agilent 1260 Infinity high-performance liquid chromatograph as described previously[37]. For *L. lactis* competition experiments, colony forming units were measured on GM17 (NZ9000 + NZ9010) or GM17 + 5ug/mL erythromycin (NZ9010 only) plates. Microscopy analysis was performed on a Leica DM4000B-M fluorescence microscope using standard brightfield and FITC filter settings.

### Double emulsion generation

The Dropception setup consists of four syringe pumps for the carrier phases (Harvard Apparatus PicoPlus Elite), a stereoscope (Amscope), a high-speed camera (ASI 174MM, ZWO), and a desktop computer (HP). Consumables included PE-2 tubing, eppendorf or anaerobic vials for droplet collection, media components, and HFE7500 oil + 2.2% Ionic Krytox (FSH, Miller-Stephenson)[38]. The cell and inner phases for the aqueous droplet cores typically contained basal bacteria growth medium (LB, M9, CDM, mBHI), 0.5 % BSA, and any indicated catabolic substrates or dyes. 10% Optiprep (Sigma) was added to the cell phase as a density modifier to ensure equal relative flow rates between the inner and cell phases during live DE generation. Cells were diluted to an OD_600_ = 0.05 in the cell carrier phase for single-cell loading (~20% of DEs contain a single cell, ~2% of DEs contain 2 cells) based on a Poisson distribution. The outer aqueous carrier phases contained matching bacterial growth media to the aqueous core, 2% Pluronix F68 (Kolliphor P188, Sigma), and 1% Tween-20 (Sigma). For anaerobic enrichments, all components were assembled inside an anaerobic chamber (COY), and all consumables were left in the anaerobic chamber to remove excess oxygen at least 3 days prior to use. Dropception device master molds for 30 and 45 μm DEs were designed in AutoCAD 2019 and fabricated via multilayer photolithography in a clean room as described previously[31]. Dropception PDMS devices were made through standard one-layer soft lithography on a standard laboratory benchtop as described previously[31]. Immediately prior to use, the outer and outlet channels of the PDMS devices were selectively O_2_ plasma treated for 12 minutes at 150 W in a Harrick PDC-001 plasma cleaner, flushed with PBS + 2% Pluronic F68, taped with scotch tape, and transferred into the anaerobic chamber or aerobic setup. The four phases (oil, cell, inner, outer) were loaded into syringes (PlastiPak, BD) and connected to the device via PE/2 tubing (Scientific Commodities). Typical flow rates were 300:100:105:6000 (oil : cell : inner : outer) μL h^−1^ for dual-inlet 45 μm devices and 275:85:2500 (oil : inner : outer) μL h^−1^ for single-inlet 30 μm devices. Live DE generation was monitored using the stereoscope and high-speed camera inside the anaerobic chamber.

### DE-FACS

DE-FACS was performed as previously described on a Sony SH800 [27, 31]. Briefly, 50 μL of DEs were diluted into 500 μL of FACS diluent buffer (PBS + 1% Tween-20) in a 5 mL 12 × 75 mm round bottom FACS tube (BD Biosciences). After standard autocalibration on the Sony SH800, the DEs were gently resuspended prior to loading, and the droplets were analyzed on a 130 μm microfluidic chip using a standard 408 nm laser configuration. DEs appear on the SH800 after 2 – 3 minutes within specific FSC-H and FSF-W gates, followed by subsequent gating on FITC fluorescence when indicated for sorting (yield mode). Drop delay adjustments for sorting were manually calibrated as described previously[31]; optimal drop delay settings typically matched those estimated by the autocalibrated Sony SH800 settings to achieve >50% DE recovery in 96 well plates for 30 μm DEs. Event rates for FACS analysis were kept below 1000 events/s and sorting rates were maintained under 50 events/s. Gain settings for DE-FACS were described previously[31], with the exception of the FITC gain, which was set to 32% or 40% for *E. coli* GFP or SYTO-stained cells, respectively. Sorted DEs were deposited into either FACS tubes or 96 well plates pre-loaded with 100 μL of osmolarically-balanced outer solution based on inner core media components. Sorted DE populations were imaged on a Leica DM4000B-M fluorescence microscope and a Leica DM E brightfield microscope.

### Mathematical modeling

The mathematical models used to simulate competitive outcomes of rate vs yield specialists was based on previous Monod models [39, 40], except the dilution term was removed to reflect growth in fed-batch cultures and the growth kinetic parameters were based on experimentally-determined monoculture growth trends collected previously [24] and within this study. Differential equations used in the basal model and extended discussions are included in the supplemental data.

### MATLAB code

The MATLAB program was customized to analyze Leica DM4000B-M fluorescence microscope images (TIFF format) with the standard brightfield and FITC filter settings overlayed (F1C). MATLAB first processed each image by sharpening droplet edges, identifying each droplet’s coordinates via the *findcircles* function, and cropping around each droplet. Then, it overlayed a white, circular mask sized with each droplet’s radii so that only the pixels inside the droplet were included in the image crop. The radii of the white masks were adjusted to remove the excess droplet edge boundaries (SF5). Then the code summed all pixel intensities above an experimentally determined threshold (data not shown) inside the droplet interior. This threshold was implemented to remove excess black pixels so that larger droplets did not return higher summed pixel intensities. The program ultimately returned a table with all indexed droplets and their respective fluorescence sums, which was then analyzed to relate number of cells to fluorescence.

### 16S sequencing

16S sequencing was performed and analyzed by the ZymoBIOMICS Service (Zymo Research, Irvine, CA). Zymo Research performed library preparation, post-library QC, sequencing using a Illumina MiSeq Platform, and bioinformatics analysis using the Dada2 and QIIME pipeline. Raw reads and metadata were submitted to SRA under BioProject PRJNA852267.

## Results

### Development of GrowMiDE platform for microbial enrichments

To encapsulate single microbial cells for parallel high-throughput culturing in DEs, we based GrowMiDE on the Dropception[31] microfluidic platform, which consists of a simple one-step microfluidic device, syringe pumps for the carrier phases, a high-speed camera, and a stereoscope (**Fig. 1A, Fig. S4**). Monodisperse DEs of 30 or 45 μm in total diameter, respectively dependent on device geometry, were generated at ~1 kHz containing: HFE7500 + 2.2% ionic Krytox[38] as the oil phase, basal growth medium + 2% pluronix F68 + 1% Tween-20 as the outer phase, and PBS or basal growth medium + 0.5% BSA + catabolic substrates as the cell and inner phases, respectively (**Fig. 1A,B**). After generating DEs within the microfluidic device, we collected them in bulk for downstream incubation or analysis (**Fig. 1B**). To ensure DEs contained only a single cell after stochastic loading, we operated within a Poisson regime in which 80% of DEs were empty and 98% of the DEs containing cells contained only a single cell (OD_600_ 0.05 in cell carrier phase). Collecting DEs for 4 hours yielded ~48 million parallel DE microreactor cultures containing single cells, allowing for an increased encapsulation of low-abundance species.

To determine if single cells encapsulated in DEs would grow, all DEs were incubated in bulk and microbial growth in droplets was assessed by brightfield or fluorescence microscopy (**Fig. 1C,D**). Overnight incubation of DEs containing single *Escherichia coli* cells constitutively expressing GFP resulted in DEs containing 30-100 cells (~5-7 generations), indicating robust growth in droplets uninhibited by DE components (**Fig. 1C**). GrowMiDE is compatible with both aerobic and anaerobic microbial growth; the Dropception setup is fully operable within an anaerobic glove box (**Fig. S4**). To demonstrate these capabilities, we successfully encapsulated and grew several facultative and strictly anaerobic microbes including *E. coli* MG1655 (performing mixed acid fermentation), *Lactococcus lactis spp cremoris* NZ9000 (lactic acid-producing fermentation), *Desulfovibrio ferrophilus* IS5 (sulfate reduction) and the thermophile *Thermoanaerobacter kivui* (acetogenesis at 65°C) (**Fig. 1D**). These results establish that DEs are a suitable high-throughput platform to cultivate diverse microbial species.

To quantify growth in DEs, we developed a custom MATLAB script to automatically detect per-droplet intensities from fluorescence microscopy images (**Fig. S5, S6**). Using GFP-expressing *E. coli* and this automated script, we analyzed growth in DEs containing either glucose or acetate in the inner phase as the sole catabolic substrates (**Fig. S7A**). Net *E. coli*-GFP growth in DEs matched batch cultures (**Fig. S7B,C,D**), indicating that *E. coli* growth in GrowMiDE and batch cultures are comparable. Together, these data demonstrate that the GrowMiDE platform can be used to cultivate and quantify growth of diverse microbial species in high-throughput.

### GrowMiDE enriches for microbial communities distinct from traditional batch cultivation

We next used the GrowMiDE platform to cultivate cells from a mixed microbial community. We focused specifically on the human gut microbiome due to knowledge of major metabolic groups of microorganisms[41–44] and established media conditions that can cultivate many different taxa[33, 45]. Specifically, we tested whether GrowMiDE cultivation enriched for novel taxa by comparing microbial enrichments performed in either batch cultures or DEs (**Fig. 2A**) (**Table S2**). 16S rRNA gene sequencing revealed a clearly distinct microbial community composition in GrowMiDE enrichments relative to batch-grown cells, even in the same basal medium (mBHI) (**Fig. 2B**). While the absolute number of recovered species did not differ significantly between batch and GrowMiDE enrichments (**Fig. 2C**), the unique identities of enriched ASVs were significantly different (**Fig. 2D**). Interestingly, GrowMiDE uniquely yielded significant growth (6.0 ± 4.0%, SD, 4/6 replicates) of the gut hydrogenotrophic methanogen *Methanobrevibacter smithii*, enriched from ~0.4% in the corresponding input stool community (**Fig. 2B, Fig. S8A,C**). The most surprising increase of a single taxa in GrowMiDE enrichments was the enrichment of the Negativicutes *Phascolarctobacterium faecium* (17.8 ± 9.8%, SD, 6/6 replicates), a 22-fold increase from the relative abundance in the input stool community (0.8 ± 0.2%, SD, n=5). (**Fig. 2B, Fig. S8B,D**). Negativicutes are a poorly understood phylogenetic group with a unique cell envelope structure, and they are thought to play a role in production of critical short chain fatty acids (SCFA) in the gut microbiome[46, 47]. The *in situ* metabolism of *P. faecium* is unknown, but its primary catabolism is thought to be secondary fermentation (succinate to proprionate)[48]. *P. faecium* has been suggested to participate in vitamin B_12_ cross-feeding with *Bacteroidetes*[49]. Strikingly, *P. faecium* is present in over 67% of human stool samples[46], however, it is difficult to isolate and, therefore, was featured on the Human Microbiome Project’s “Most Wanted” list[33].

**Figure 2.**
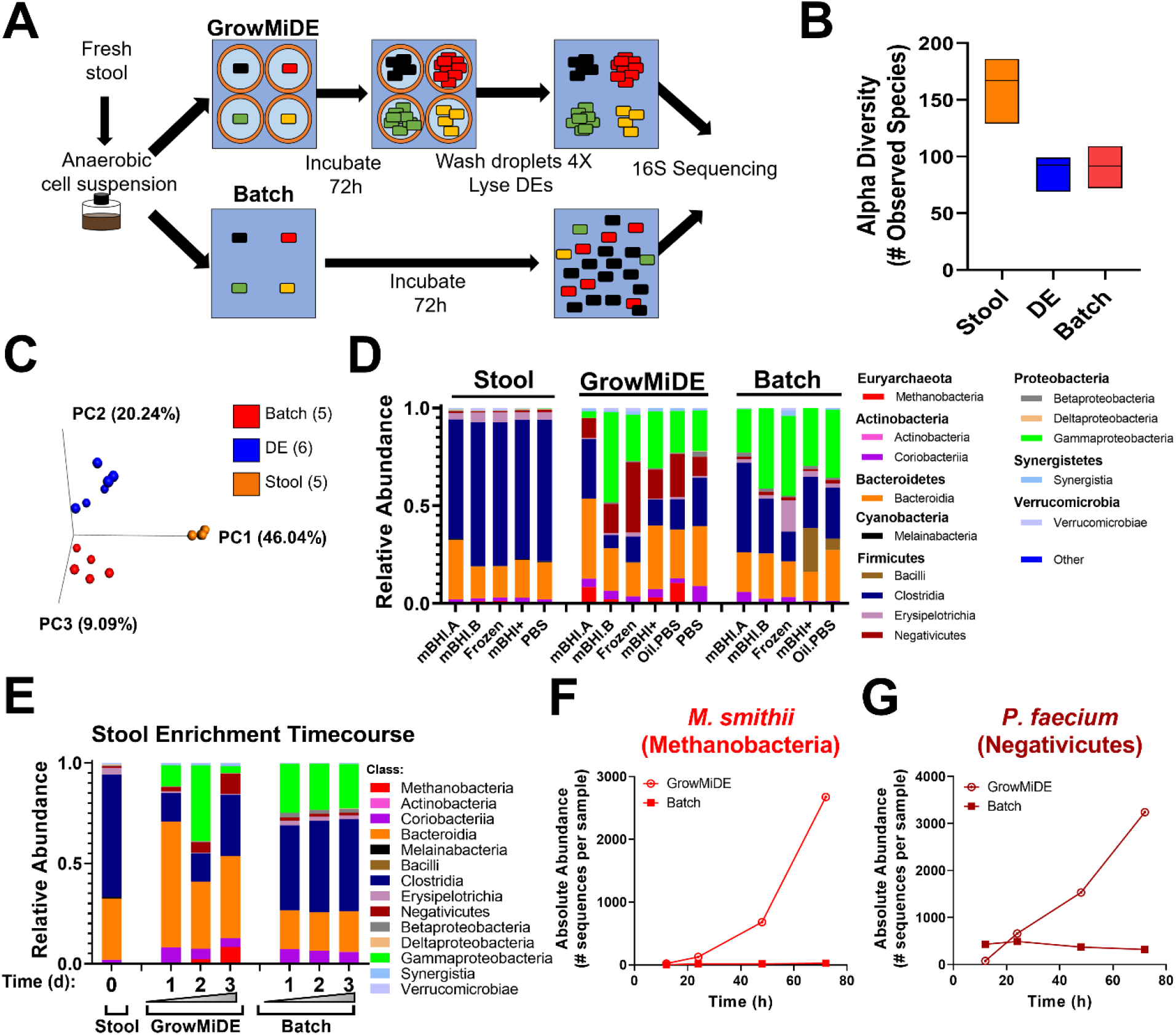
GrowMiDE enrichment of distinct microbial communities from human gut microbiota. (**A**) Schematic overview of stool enrichments in DEs vs batch enrichments in mBHI. (**B**) Alpha diversity from input stool, GrowMiDE, and batch enrichments based on total unique ASVs. Floating bar plots represent the mean and range, n=5-6. (**C**) Beta diversity from input stool, GrowMiDE, and batch enrichments based on type of unique ASVs. (**D**) Relative 16S rRNA gene abundances from input stool samples, GrowMiDE enrichments, and batch enrichments from stool cell suspensions. (**E**) Relative 16S rRNA gene abundances from a timecourse of stool enrichments in DE vs batch cultures sacrificed at 12, 24, 48, and 72 hours, and corresponding absolute 16S rRNA gene abundances of (**F**) *M. smithii* (class: Methanobacteria) and (**G**) *P. faecium* (class: Negativicutes).

In batch cultures, the final microbial composition was mostly established within 12 hours, consistent with growth dominated by the fastest species in an artificial laboratory environment (eg *E. coli* and *Enterococcus spp*,) (**Fig. 2E**). However, in GrowMiDE enrichments we observed gradual enrichment and increased gene copies of *M. smithii* and *P. faecium* over time (**Fig. 2E-G**). The appearance of these taxa at later timepoints and their enrichment across time support the hypothesis that the GrowMiDE platform decreased the bias against slower-growing species and facilitated enrichment of novel physiologies, even with non-selective medium.

### GrowMiDE favors growth yield specialists over growth rate specialists

We hypothesized that stool GrowMiDE cultures enriched for slower-growing taxa by minimizing competition for shared nutrients between droplets (i.e. nutrient privatization). Nutrient privatization through spatial structuring promotes diversity in communities, including microbes that prioritize growth yield over rate that would otherwise be lost in laboratory enrichments (**Fig. S1C**). Trade-offs between rate and yield-specialists has been well-documented in *Lactococcus lactis* strains[24, 34](**Fig. 3A**). Under high glucose flux conditions, wild type *L. lactis* (NZ9000) ferments 1 mol glucose to 2 mol lactate by homofermentative lactic acid fermentation (**Fig. S2A**), resulting in a net yield of 2 mol ATP per mol glucose. Under conditions of low glucose flux, fermentation shifts towards formation of ethanol and acetate (**Fig. S2B**), resulting in a total net gain of 3 mol ATP per mol glucose. This latter fermentative metabolism is locked in a *L. lactis* lactate dehydrogenase deletion mutant (∆*ldhA*, NZ9010). Although the ∆*ldhA* strain has a higher growth yield, it comes at a trade-off in a reduction of the maximum growth rate (**Fig. 2B**). In elegant single emulsion experiments, encapsulation of WT and ∆*ldhA* strains resulted in enrichment of the efficient, but slower-growing ∆*ldhA* strain across transfers[24].

**Figure 3.**
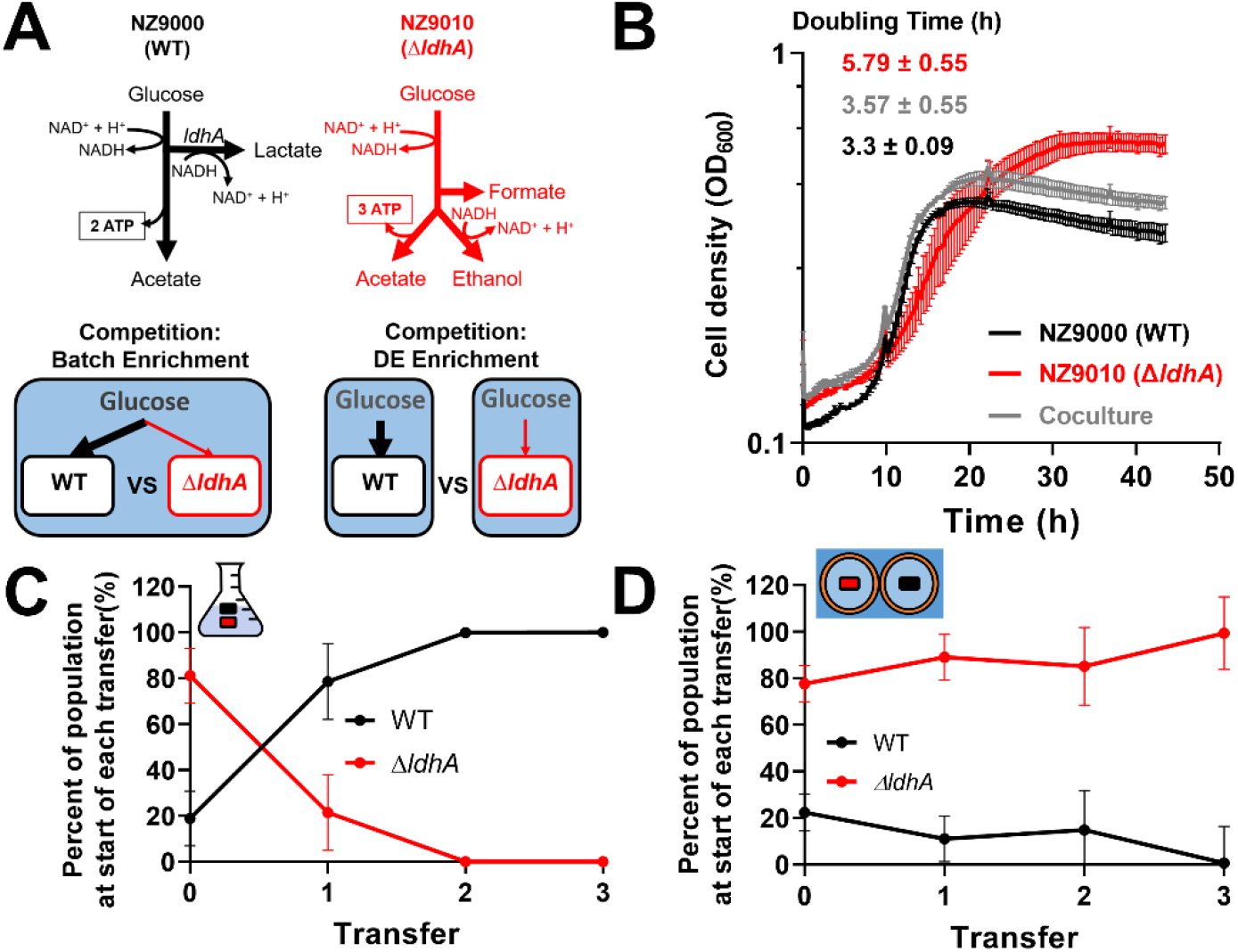
GrowMiDE can maintain and enrich growth yield specialists through nutrient privatization. (**A**) Fermentation pathways of *L. lactis* WT (black) and ∆*ldhA* (red) strains in mixed batch cultures or GrowMiDE enrichments. (**B**) Growth of *L. lactis* monocultures and cocultures (1:1 starting ratio) on CDM + 25 mM glucose (n = 3, biological replicates, error bars indicate SEM). (**C**) Frequencies of populations across transfers in mixed batch cultures of *L. lactis* strains on 50 mM glucose. Each transfer received a 1% inoculum into fresh CDM + 50 mM glucose, and CFUs for each strain were determined from the grown community at each transfer (n = 3, biological replicates, error bars indicate SEM). (**D**) Cell densities across transfers in GrowMiDE enrichments of *L. lactis* strains on 50 mM glucose. Initial mixed culture contained ~80% ∆*ldhA* as determined by CFUs/mL on CDM + Ery^5^ plates. At the end of each transfer, the DEs were disrupted in bulk by incubating with 1H,1H,2H,2H-perfluoro-1-octanol (PFO), diluted to an OD = 0.05 to achieve single-cell loading, and cells were re-packaged into DEs for the next transfer. (n = 3, technical replicates, error bars indicate SEM). CFUs for each strain were determined from the grown community at each transfer to determine relative cell densities, and the initial mixed culture contained ~80% ∆*ldhA* prior to splitting into batch or DE competition experiments. All *L. lactis* transfers were incubated for 48 hours at 30°C.

To test whether DEs similarly maintained populations with higher growth yield, we competed *L. lactis* WT and ∆*ldhA* strains in batch vs GrowMiDE enrichments over serial transfers. Growth rates of batch cocultures containing a 1:1 mixture of both strains resembled the faster WT strain (**Fig. 3B**), which was consistent with WT constituting over 80% of the culture after transfer 0 (**Fig. 3C**). As expected, even when the ∆*ldhA* strain was inoculated at a high frequency of 80%, it was outcompeted to less than 0.003% by WT within 2 transfers in batch culture (**Fig. 3C**)[24]. In contrast, the slower ∆*ldhA* population was maintained and even dominated WT in GrowMiDE enrichments across transfers (99.33% ∆*ldhA* by transfer 4) (**Fig. 3D**), showing that GrowMiDE can retain or even enrich slower-growing species from mixed communities through nutrient privatization.

### GrowMiDE + DE-FACS as a high-throughput enrichment and isolation tool

A useful feature of DE platforms is direct compatibility with traditional microscopy or FACS (DE-FACS) (**Fig. 4A**) [31, 32, 50, 51]. Developing a platform that facilitates high-throughput encapsulation and the ability to selectively isolate droplets of interest has broad applications including improved genome recovery[52], antimicrobial sensitivity assays[16, 53], and directed evolution studies[54]. To test whether our cell-containing DEs could be accurately distinguished and isolated from empty DEs via fluorescence during FACS, we encapsulated and grew *E. coli* cells expressing GFP in DEs and then attempted to isolate only cell-containing droplets via FACs. Distinct FACS populations of empty and cell-laden DEs were observed by FACS, and fluorescent microscopy measurements confirmed that 97.5% of the sorted positive population contained intact DEs loaded with cells (**Fig. S9A**). We also assessed whether DE-FACS can differentiate between empty and microbe-containing DEs by adding a fluorescent dye (SYTO_bc_) during cell encapsulation. We identified FACS gates that distinguished empty DEs from DEs containing *E. coli* MG1655 loaded with 2.5 μM SYTO_bc_ grown for 24 hours (**Fig. S9B**). Within the SYTO^+^ DE population, 92.2% contained cells as confirmed by brightfield microscopy based on detecting cell motility. Longer incubation times decreased fluorescence signal; however, SYTO^+^ DE populations were still detectable at 48 hours (**Fig. 4C**). Relative to the input GrowMiDE enrichment that contained ~80% empty DEs, DE-FACS resulted in a 443- and 419-fold enrichment for DEs containing *E.coli*-GFP and *E.coli* + SYTO_bc_ DEs, respectively. These results demonstrate the application of DE-FACS to sort for DE droplets containing grown bacterial cells.

**Figure 4.**
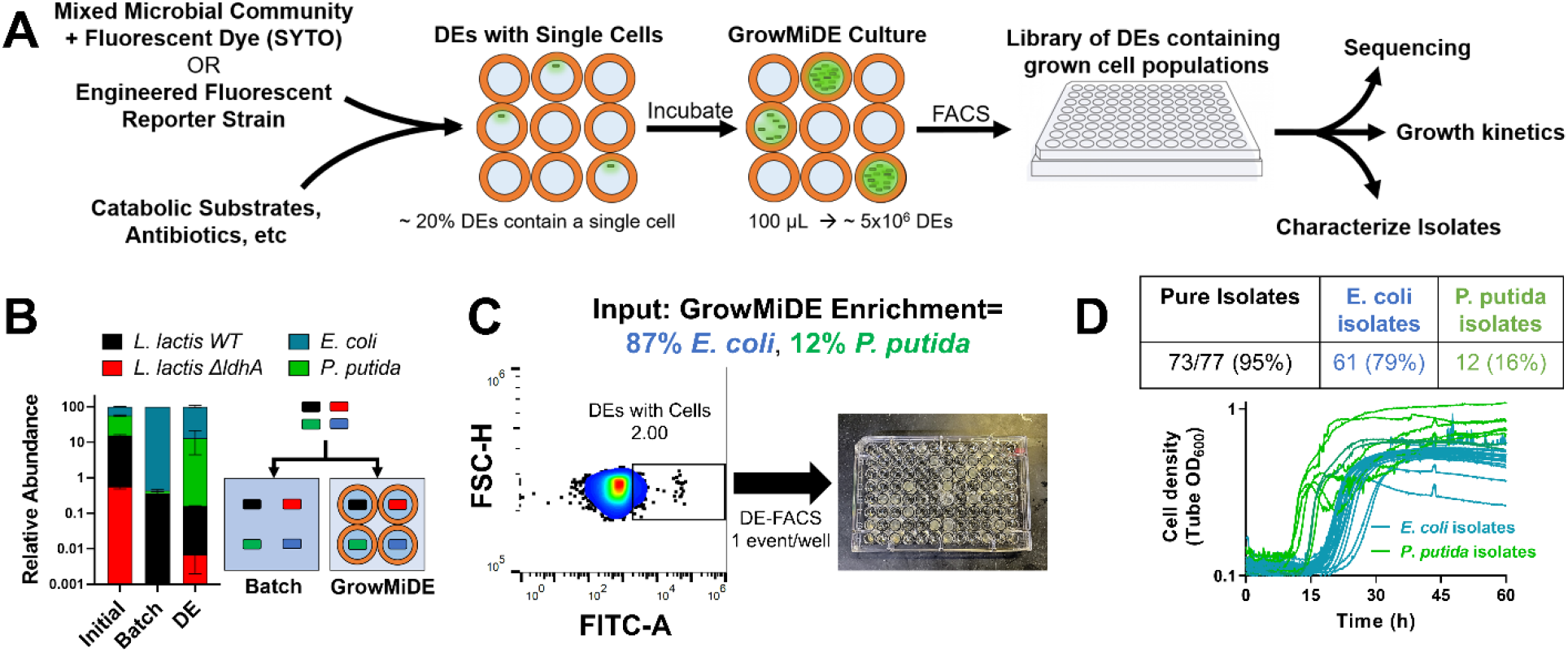
GrowMiDE + DE-FACS as a novel microbial enrichment and isolation platform. (**A**) Overview and potential applications for microbial enrichments using GrowMiDE followed by DE-FACS to obtain isolates. (**B**) Relative ratios of 4 glucose-catabolizing strains in a mock community enriched in batch cultures or the GrowMiDE platform on CDM + 25 mM glucose for 48 hours. n=3. Error bars indicate SEM. (**C**) FACS profiles of 30 μm DEs containing the mock community encapsulated with 2.5 μM SYTO9 after 48 hours and subsequently sorted into individual wells on a 96-well plate. Plate wells were preloaded with growth medium and 10uL of PFO to disrupt DEs. 767 events were gated within the top 2% of the SYTO^+^ DE population (38321 events) and 192 were sorted onto plates to assess for growth. (**D**) Growth curves of 34 representative output isolates downstream of DE-FACS. All growth assays were performed at 30°C.

A major barrier in single emulsion microfluidics and traditional single-cell FACS sorting is the difficulty in isolating live cells for downstream phenotypic analysis. While single-cell genomes can easily be recovered in downstream processes even if cells die, preserving cell viability in sorted populations is essential for downstream characterization of microbial physiology. High-throughput microbial cultivation using GrowMiDE and isolations using DE-FACS therefore has the potential to be a powerful approach for screens, enrichments, and isolations (**Fig 4A**). As a proof of concept, we demonstrated the application of GrowMiDE + DE-FACS in a synthetic glucose-catabolizing community containing *E. coli*, *Pseudomonas putida*, *L. lactis* WT, and *L. lactis* ∆*ldhA*. The synthetic community was chosen due to i) compatibility of all species in a defined medium, ii) a comparable catabolic usage of glucose, and iii) the ability to use selective plating to phenotypically assess species identities from mixed cultures. *E. coli* dominated both batch and GrowMiDE enrichments due to higher relative growth rate and yields in pure cultures (**Fig. S10**); however, only GrowMiDE cultivation preserved all 4 strains in the synthetic community (**Fig. 4B**, **Fig. S11**). Illustrating how even minor decreases in maximal growth rates can drive species lost in batch cultures, *P. putida* was frequently lost in batch culture competitions after 48 hours despite having the second highest growth rate and yield in monocultures (**Fig. S10**). For GrowMiDE enrichments, we sorted individual DEs from the SYTO^+^ DE population (**Fig. 4C**) into 96 well plates, released encapsulated clonal populations from DEs with PFO, and collected growth measurements of recovered isolates. Here, we used 48 hour incubations despite decreased stained bacterial intensities and resulting broader FACS gates to provide sufficient time for all strains to grow in DEs (**Fig. 4C, Fig. S10**). Our approach included dimly-fluorescent DEs to limit bias in staining efficiency between different species with the trade-off of sorting a higher fraction of empty DEs (**Fig. 4C**). Across two 96-well plate arrays, 77 wells contained bacteria recovered from sorted DEs, 73 (95%) of which were confirmed to be pure cultures by selective plating for all 4 input species (**Fig. 4D**). Consistent with the 2 dominant species abundances from GrowMiDE enrichments (87% *E. coli* : 12% *P. putida*), 79% of the recovered pure cultures were *E. coli* and 16% were *P. putida*. Together, these results demonstrate the feasibility of microbial enrichments and isolations using GrowMiDE + DE-FACS.

## Discussion

Here, we demonstrated a novel application of DE technology to cultivate microbes and perform high-throughput enrichments for slower-growing microbes in both defined and undefined communities. We demonstrated that many metabolically diverse microbial species can grow in GrowMiDE, including strict anaerobes (**Fig. 1**). Moreover, GrowMiDE can also facilitate enrichment for microbes pursuing a growth yield over growth rate strategy (**Fig. 3D**). GrowMiDE enrichments uniquely yielded microbial species that were never observed in batch culture using the same enrichment medium (**Figs. 2,4**). Finally, we demonstrated the application of GrowMiDE and DE-FACS as a platform to obtain isolates for downstream characterization and physiological studies (**Fig. 4**).

In our study, nutrient privatization allowed slow-growing species to escape competition [1, 2]. Although DEs spatially separate cells and large macromolecules in separate microreactors, the inner aqueous core is only separated from the environment by a thin oil shell. Dependent on the local surfactants and blocking agents used in the oil layer, DEs can therefore be selectively permeable to small molecules including oxygen, salts, and some dyes[28–30, 55]. This effect is reported to be tunable dependent on surfactant properties and concentrations[55]. Other rheological studies in DEs have also shown that rhodamine A, BSA conjugates, and anhydrotetracycline can traffic across the oil shell in a pH-mediated delivery system in some DE formulations[29]. The molecular mechanism of crossover is unknown, although facilitated diffusion and spontaneous emulsification have been hypothesized[29, 56].

While it is possible that low levels of carbon source crossover occurred within our GrowMiDE enrichments, the levels of glucose retained within the DEs remained sufficient to favor growth of slow-growing *L. lactis* ∆*ldhA* mutants (**Fig. 3D**). Degrees of nutrient privatization can be dynamic; even partial privatization, in which limited competition exists, can be sufficient to preserve distinct populations[57–59]. Another possibility is that low permeability of DEs for certain compounds could have facilitated the growth of *M. smithii* and/or *P. faecium*. *P. faecium* and *M. smithii* have been indicated to be fully dependent on cross-fed metabolites from the gut community, succinate and H_2_ respectively; however, neither compound was added to the GrowMiDE enrichments. H_2_ is produced during fermentation by many co-enriched gut species and should readily diffuse across the HFE7500 oil. For *P. faecium* enrichments, it is unclear how picomolar concentrations of succinate crossover from a subsection of a mixed community would promote a significant enrichment. Full genome sequence analysis of the two existing isolated *P. faecium* strains suggests that fermentation of succinate to proprionate is their only primary catabolism[60]. However, it is possible that the GrowMiDE enrichment facilitated the use of other catabolic substrates that were available in our enrichments, either in the basal mBHI medium or produced by another species. Control batch culture enrichments supplemented with DE components did not enrich for *P. faecium* (data not shown), ruling out the possibility of catabolism of oil or DE surfactants.

Recent advances in microfluidic technology to expand the number of cultured species include custom devices to isolate and incubate single cells (iChip, SlipChip)[61–63] and culturomics, in which bacterial colonies are arrayed across media conditions on plates and identified using MALDI-TOF[45]. However, these technologies are limited in throughput and require specialized, expensive equipment to analyze and sort grown cultures. The GrowMiDE platform is highly amenable to diverse applications and requires only simple instrumentation, all of which can be assembled and operated with minimal training. In this study, we showed that GrowMiDE eliminated competition with faster growing microbes and enriched microbes from a mixed community that are categorically lost in batch cultures under the same media conditions (**Fig 4**). Therefore, developing, high-throughput enrichment and isolation methods like GrowMiDE + DE-FACS that do not favor fast-growing microorganisms might serve as a novel approach to combat ‘The Great Plate Count Anomaly’.

## Acknowledgments

We thank H. Bachmann for the *L. lactis* strains NZ9000 and NZ9010, and A. Maheshwari for the *E. coli*-GFP strain. We also thank members of the Fordyce lab Dropception team and Prof. A. Bhatt’s lab for helpful discussions on experiments. This work was supported in part by the US National Science Foundation through the Center for Deep Dark Energy Biosphere Investigations, N.I.H. 1DP2 GM123641 awarded to P.M.F, and a Simons Postdoctoral Fellowship in Marine Microbial Ecology to A. L. McCully from the Simons Foundation, Division of Life Sciences, award #600755. P.M.F. is a Chan Zuckerberg Biohub Investigator.

## Data Availability Statement

All raw data and Matlab scripts are freely available to any researcher upon request. 16S sequencing data has been submitted to SRA under BioProject PRJNA852267.

