## Supporting Information for "Double emulsions as a high-throughput enrichment and isolation platform for slower-growing microbes"

**This PDF file includes:**

Supplementary text
Figures S1 to S11
Tables S1 to S2
SI References

#### Extended Methods for GrowMiDE Enrichments

*E. coli*: An overnight culture of MG1655 or Ec-GFP was washed with M9 medium and used as the inoculum for all *E. coli* double emulsion experiments. Cells were diluted to an OD of 0.05 in medium (LB, M9, or mBHI) + 0.05% BSA + 10% Optiprep and loaded into a 1 mL syringe for the cell carrier phase in dual inlet 45 µm DE devices. The inner phase consisted of the same basal medium, indicated catabolic substrates (glucose, acetate,), and 0.5% BSA as a stabilizing agent.

*Lactococcus lactis: S*tationary phase cultures of NZ9000 or NZ9010 were washed with CDM medium, combined into a single tube and diluted as the inoculum for all *L. lactis* competition experiments. Cell densities were taken at the start of the experiment to determine relative CFUs/mL by plating on GM17 (NZ9000 and NZ9010) and GM17 + Ery$^{5}$ (NZ9010 only). For batch competition experiments, the washed mixture of *L. lactis* populations were diluted to an OD of 0.005 in CDM supplemented with 25 mM glucose. After 48 hours of static growth at 30°C, 1% of the cultures were transferred to fresh 10 mL CDM + glucose in biological triplicates.  For DE competition experiments, *L. lactis* cells were diluted to an OD of 0.05 in CDM medium + 0.05% BSA + 10% Optiprep and loaded into a 1 mL syringe for the cell carrier phase in dual inlet 45 µm DE devices. The inner phase contained CDM medium, 50 mM glucose, and 0.5% BSA as a stabilizing agent. The outer phase contained CDM medium, 2% Pluronix F68, and 1% Tween-20. After 48 hours of static growth at 30°C, the DEs were washed 3 times in an equivalent volume of freshly prepared outer solution (CDM + 2% Pluronix F68 + 1% Tween-20) to remove escaped cells. The DEs were then gently lysed through addition of a 1:1 mixture of 1H,1H,2H,2H-Perfluoro-1-octanol (PFO) and gently flicking for 15 minutes or until no intact DEs remained. The top aqueous layer containing the grown *L. lactis* populations was removed and used as the inoculum for the next transfer in DEs in technical triplicates. Cell densities of each *L. lactis* population were monitored at the start of each transfer by plating for CFUs/mL.

Stool samples: Fresh stool from a healthy donor was immediately transferred into anaerobic conditions using a GasPak jar and stored at 4°C for 1-12 hours. Fresh stool samples were then transferred into an anaerobic glove box to extract cell suspensions. Briefly, a ratio of 5 mL PBS to 1 g of fresh stool was added and stirred at max speed for 15 min or until homogenous. The resulting liquid suspension was filtered through a coffee filter to remove large particles and centrifuged at low speed to settle smaller particles (6000 rpm for 5 min). The resulting cell suspension was removed from larger particulates, diluted into ice-cold anaerobic mBHI medium at an OD ~ 1 prior to enrichments, and a sample was immediately frozen at -80°C. Aliquots were frozen in anaerobic vials with equal volumes of 50% glycerol and stored as frozen samples. 5 biological replicates of input stool samples from the same healthy donor collected over a period of 44 days were used for DE enrichments. For batch enrichments, a 1% inoculum was transferred into 10 mL of anaerobic mBHI medium and incubated at 37°C for 72 hours. For DE stool enrichments, cell suspensions were diluted to an OD = 0.05 in the cell carrier phase consisting of medium (PBS, mBHI, or $mBHI^+$), 0.05% BSA, and 10% Optiprep. Inner phase consisted of medium (mBHI or $mBHI^+$) + 0.5% BSA, and the outer phase contained mBHI + 2% Pluronic F68 + 1% Tween-20. All DE stool enrichments were collected in a 20 mL serum vial in the anaerobic chamber, flushed with $N_2$, and incubated at 37°C for 72 hours. To harvest gDNA from stool DE enrichments, the DEs were washed 4 times with 10 mL of freshly prepared outer solution (mBHI + 2% Pluronix F68 + 1% Tween-20) and lysed using PFO as described above. The enriched community was frozen and stored at -80°C for batched gDNA harvesting. The

Powersoil kit (Qiagen) was used to extract gDNA from enriched DE populations, with the modification of an additional incubation at 65° C for 10 min prior to bead-beating to promote cell lysis. gDNA was quantified by Qubit and stored at -80°C until 16S sequencing.

Mock Community: The mock community containing *E. coli*, *Pseudomonas putida*, *L. lactis* WT, and *L. lactis* Δ*ldhA* was inoculated using washed starter monocultures of each strain grown in CDM + 50 mM glucose. Batch and GrowMiDE enrichments were performed similarly to *L. lactis* competition experiments in CDM + 25 mM glucose, except 2.5 uM SYTO$_{bc}$ was added during encapsulation. Cell densities were determined by selective and differential plating to determine ratios of *E. coli* (MacConkey), *Pseudomonas putida* (Cetrimede), *L. lactis* WT (GM17 + 40 ug/mL nalidixic acid), and *L. lactis* Δ*ldhA* (GM17 + 40 ug/mL nalidixic acid + 5ug/mL erythromycin). A 100 uL aliquot of GrowMiDE enrichments were sorted by DE-FACS (1 event/well, yield mode) using autocalibrated droplet delay settings on a Sony SH800. DEs containing cells were collected in 96 well plates containing 260 uL CDM + 50 mM glucose + 10 µL PFO. 96 well plates were incubated in a Tecan plate reader for ~65 hours at 30ºC, and wells containing growth were plated to identify isolates.

**Modeling fitness of rate vs yield specialists**

There is a huge demand for novel culture-based methods to isolate new species that continue to be overlooked in their natural communities. In addition to discovering new microbial metabolic potentials, developing approaches to reduce the bias for fast growth rates will help reveal dynamic microbial physiologies that have long been overlooked across diverse environments[1–3]. Microorganisms that prioritize growth yield over growth rate might play central roles within communities[4,5], however slower-growing species will always be outcompeted in laboratory enrichment attempts in batch culture. In a simplified two-member coculture consisting of a growth rate specialist (R) and a growth yield specialist (Y) growing on glucose (**Fig. S1A**), the outcome of a batch enrichment is entirely dependent on relative differences in growth rates (**Fig. S3A**). The rate specialist will always outcompete the yield specialist, even when the yield specialist has a physiologically impossible 100x increase in growth yield on glucose (**Fig. S3B**). Outcompetition of the slower strain can be temporarily prevented by increasing starting cell densities (**Fig. S3C**), however the relative increase required to overcome competition is dependent on actual cell densities of the fast strain (**Fig. S3D**), and will not prevent outcompetition when the mixed culture is transferred subsequently (**Fig. S1B**). Even when a mixed culture contained 99% growth yield specialist Y, the growth rate specialist R completely overtook the culture within 5 transfers in a batch culture system. The decreased competitive fitness in slower strains is due to competition for a shared nutrient pool that is equally available to both populations. However, privatization of nutrients can create more niches within a community by eliminating the bias for solely fast growth rate[6,7]. In the simplified two-member coculture model, when the R and Y populations no longer compete for the same glucose pool across multiple transfers, this results in a community composition that converges to containing 77% of the growth yield specialist, which directly reflects the difference in relative growth yields (**Fig. S1C**). These simulated results are consistent with ecological theory and empirical data which indicates that creating privatized nutrient pool can promote maintenance or even enrichment of slower, but more efficient species.

**Mathematical Model**

Rate specialist R growth rate:

$\mu R = \mu R_{Max} * (Glu/(K_g + Glu))$

Yield specialist Y growth rate:

$\mu Y = \mu Y_{Max} * (Glu/(K_g + Glu))$

Change in cell densities over time:

$dR/dt <- \mu R * R$

$dY/dt <- \mu Y * Y$

Change in extracellular metabolites over time:

$dGlu/dt <- -(\mu R * R/Y_r) - (\mu Y * Y/Y_y)$

$dLac/dt <- (R*\mu R*F_{l1}) + (Y*\mu Y*F_{l2})$

$dAce/dt <- (R*\mu R*F_{a1}) + (Y*\mu Y*F_{a2})$

$dEtOH/dt <- (R*\mu R*F_{e1}) + (Y*\mu Y*F_{e2})$

$dFor/dt <- (R*\mu R*F_{f1}) + (Y*\mu Y*F_{f2})$

where,

$\mu$ is the specific growth rate of R or Y species ($h^{-1}$)

$\mu_{Max}$ is the maximum specific growth rate of R or Y ($h^{-1}$)

$K_g$ is the half saturation constant for glucose (mM)

Glu, Lac, Ace, EtOH, and For are glucose, lactate, acetate, ethanol, and formate, respectively (mM)

R and Y are the cell densities of rate and yield specialists, respectively (cells/mL)

$Y_r$ and $Y_y$ are the cell growth yields of the rate and yield specialists, respectively, on glucose in CDM (cells/μmol glucose)

F is the fraction of glucose converted into the indicated metabolite per R or Y cell (1 and 2 respectively) based on HPLC fermentation profiles of *L. lactis* strains (μmol/cell)

### Figures and Tables

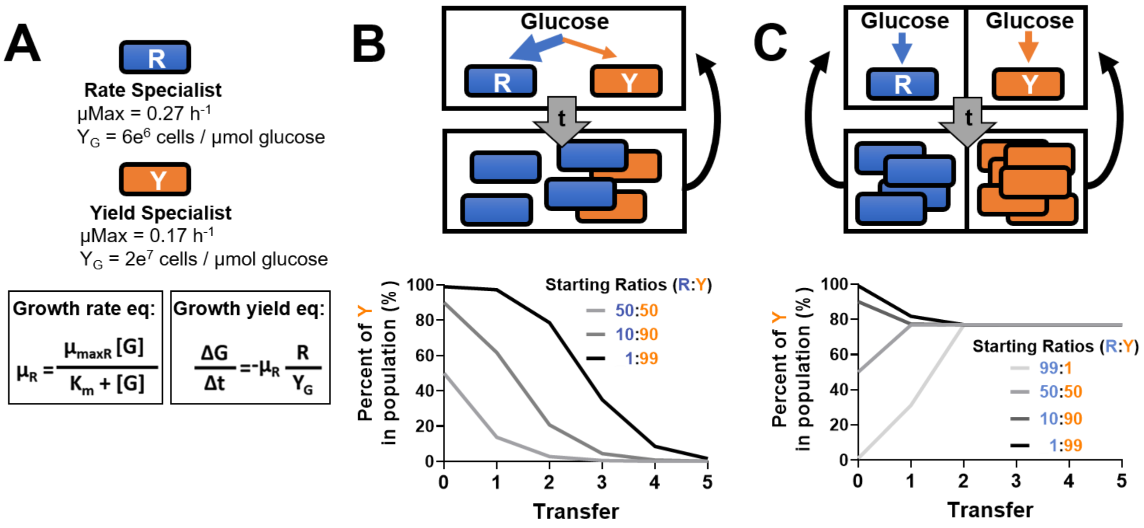


**Supplemental Figure S1. Preserved growth of slower, yield-specialist cells through nutrient privatization.** (**A**) Mathematical Monod model equations used to simulate growth of rate (R) vs yield (Y) specialists on glucose. Values for differences in maximal growth rate and growth yields are based on empirical data from *Lactococcus lactis* growth curves (**Fig. S2**). (**B**) Simulated growth of R and Y specialists in mixed batch cultures started at different starting ratios of R:Y across serial transfers. (**C**) Simulated growth of R and Y specialists separated into single-cell compartments started at different starting ratios of R:Y in compartments across serial transfers. Simulated cultures were transferred by pooling the resulting grown community and re-diluting into individual single-cell compartments. All simulated cultures were modeled for 48 hours on 25 mM glucose and 1% of the resulting community was transferred after growth.

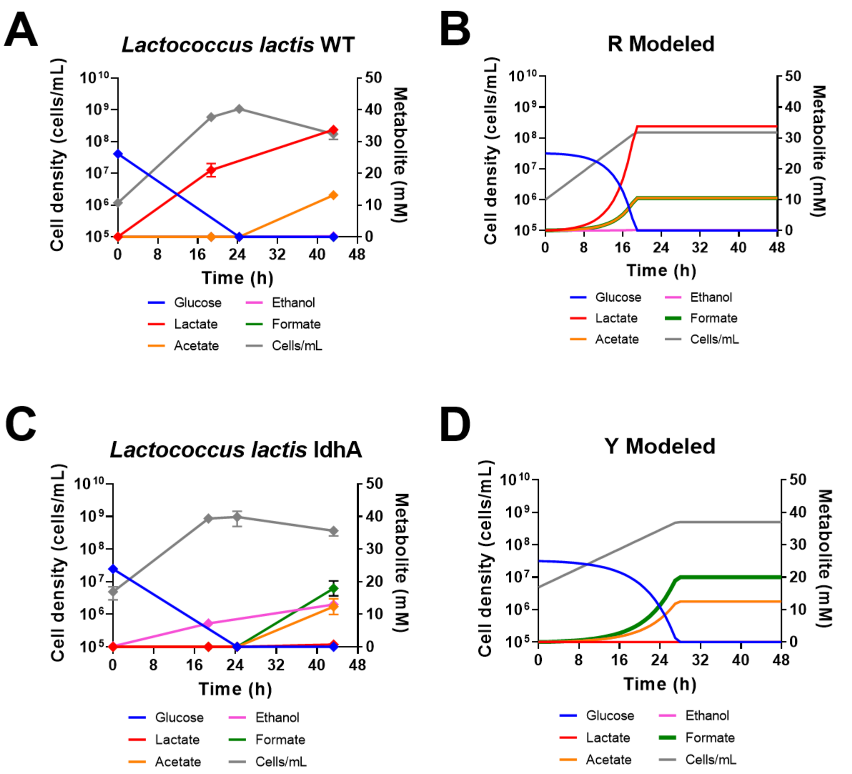


**Supplemental Figure S2. Mathematical modeling of growth rate vs growth yield specialists.** (**A**, **C**) Growth curves and fermentation profiles from *L. lactis* WT and Δ*ldhA* strains. (**B**, **D**) Fitted simulations of WT (R) and Δ*ldhA* (Y) strains in a mathematical Monod model to derive relative maximal growth rates and growth yields.

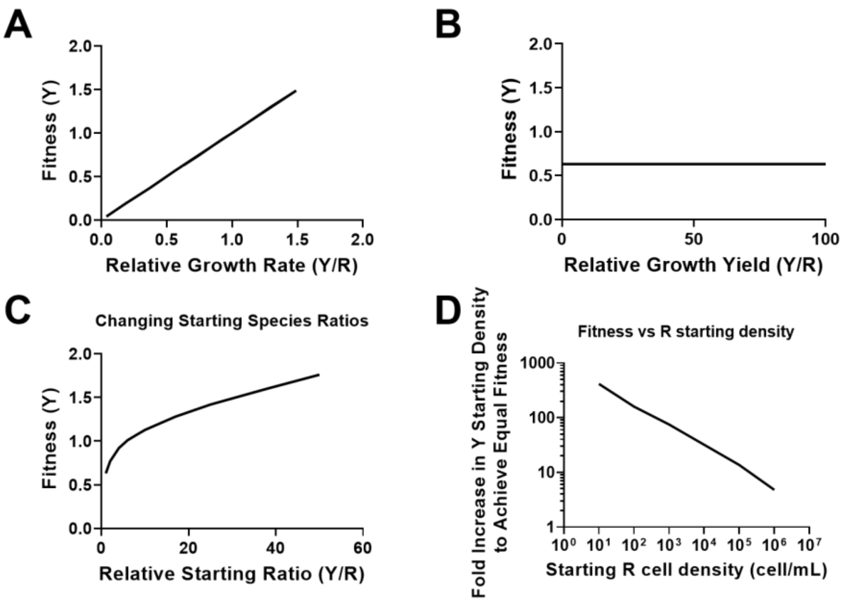


**Supplemental Figure S3. Effects of relative growth rates, growth yields, and starting cell densities on yield specialist fitness.** Fitness of yield specialists (Y) in competition with rate specialists (R) when different growth parameters are simulated including modeling relative growth rates (**A**), relative growth yields (**B**), and relative starting species ratio (**C**). (**D**) Required fold increase in Y in the initial population required to overcome outcompetition by R at different R starting cell densities.

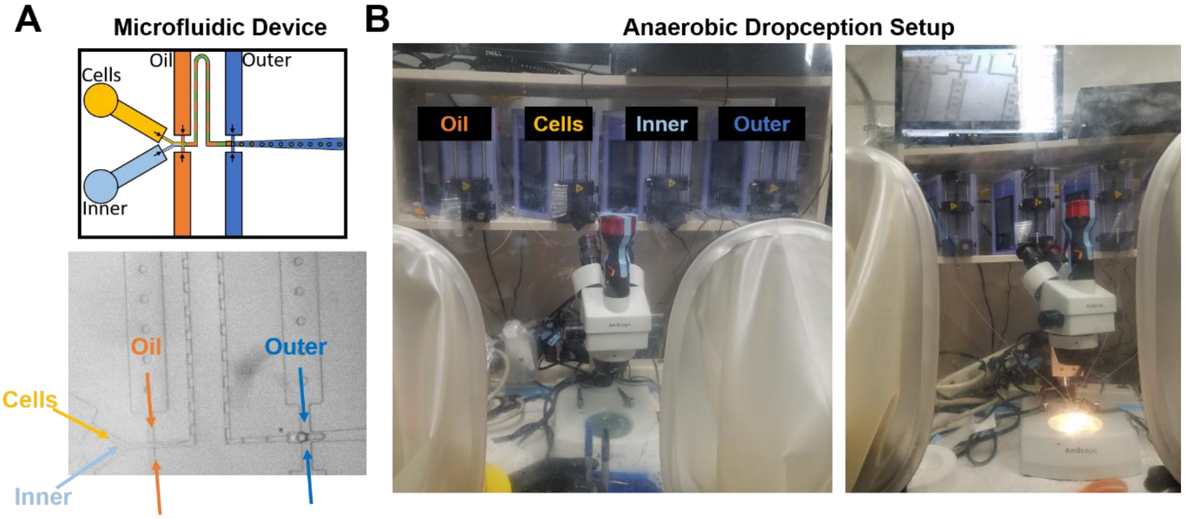


**Supplemental Figure S4. Anaerobic Dropception setup.** (**A**) Schematic (top) and labeled bright field microscopy image (bottom) of microfluidic device layout used to generate monodisperse DEs. Flow rates of carrier solutions were controlled by external programmable syringe pumps. (**B)** Photos of anaerobic Dropception setup during operation within an anaerobic chamber.

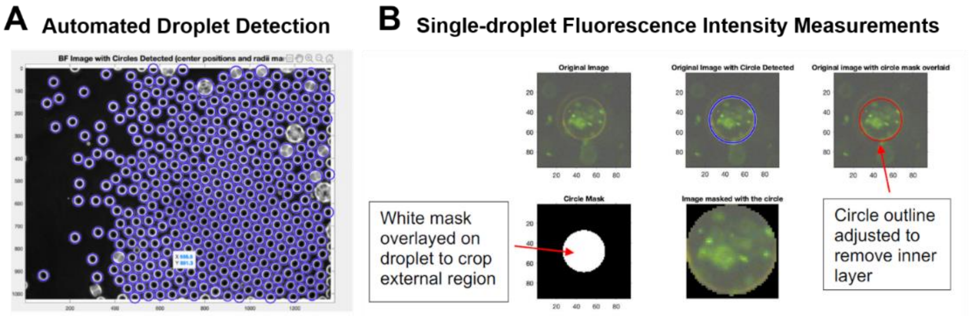


**Supplemental Figure S5. Custom MATLAB code to automate fluorescent droplet detection and quantification.** (**A**) Example brightfield microscopy image at 10X magnification showing detection of DEs from a custom MATLAB script. (**B**) Example images illustrating automated droplet detection and fluorescence quantification process: (i) droplets were detected from brightfield images using MATLAB's *findcircles* function, (ii) the identified circles were adjusted by the user to delineate margins expected for monodisperse 45 µm droplets, and (iii) a binary mask was applied to quantify the summed fluorescence intensity within the circle.

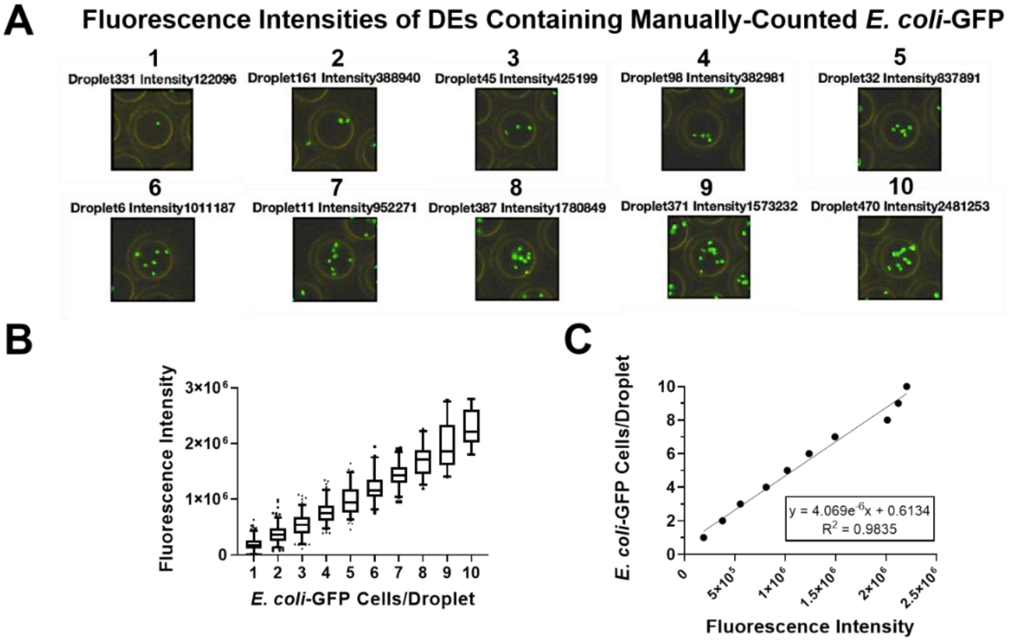


**Supplemental Figure S6. Calibrating fluorescence intensities to cell counts on a per droplet basis.** (**A**) Representative images from MATLAB showing DEs containing between 1-10 *E. coli*-GFP cells per droplet immediately after encapsulation. Cell loading was controlled by increasing cell densities in the cell carrier phase according to a Poisson distribution. (**B,C**) Manual cell counts per droplet were plotted against the summed pixel fluorescence intensities per droplet to generate a standard curve to approximate cell numbers per droplet.

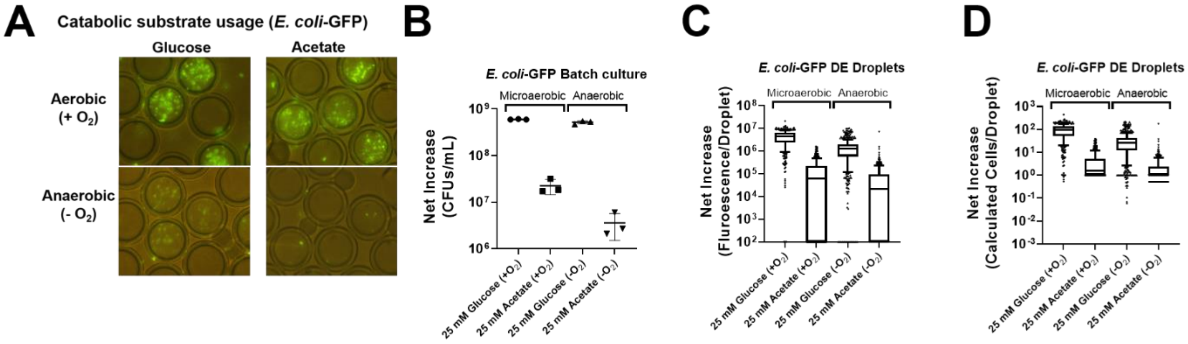


**Supplemental Figure S7. *E. coli* growth in DEs is comparable to batch culture.** Representative microscopy images (**A**) and quantification of net *E. coli*-GFP growth in batch cultures (**B**) and DEs (**C,D**) across different conditions. 25 mM glucose and acetate in the inner phase were chosen as catabolic substrates for *E. coli*-GFP during growth under anerobic or microaerobic conditions. Prior to imaging, all batch cultures and DEs were exposed to ambient air for at least 10 minutes to facilitate aerobic recovery of GFP fluorescence[8]. Error bars indicate SD, n=3. Box and whisker plots indicate 10-90$^{th}$ percentile. Net increase in *E. coli*-GFP per droplet (**D**) was calculated based on standard curves correlating manual cell counts to mean fluorescence measurements (**Fig. S7**).

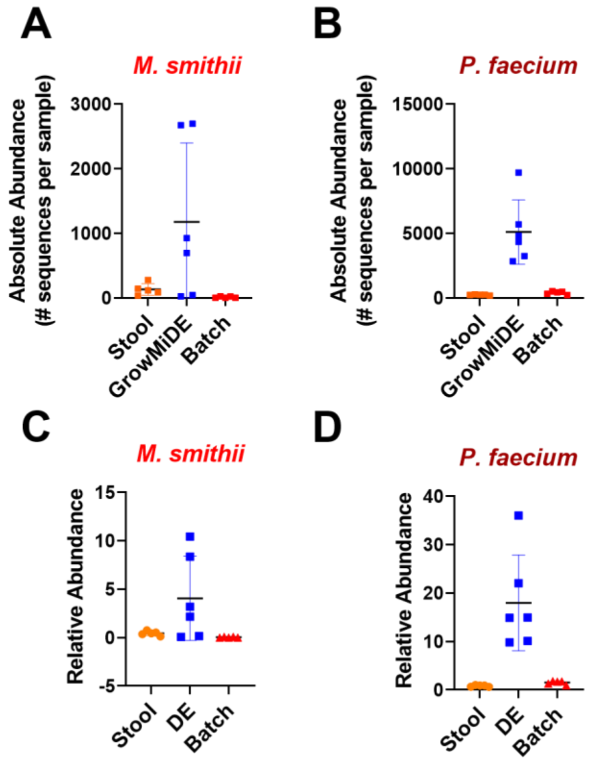


**Supplemental Figure S8. 16S rRNA gene abundances of *M. smithii* and *P. faecium* in input stool samples, GrowMiDE enrichments, and batch culture enrichments.** Absolute (**A,B**) and relative (**C,D**) abundances of 16S rRNA gene sequences belonging to *Methanobrevibacter smithii* (**A,C**) *Phascolarctobacterium faecium* (**B,D**) in input stool, GrowMiDE enrichments, and batch culture enrichments after 72 h. Error bars indicate SD, n=5-6.

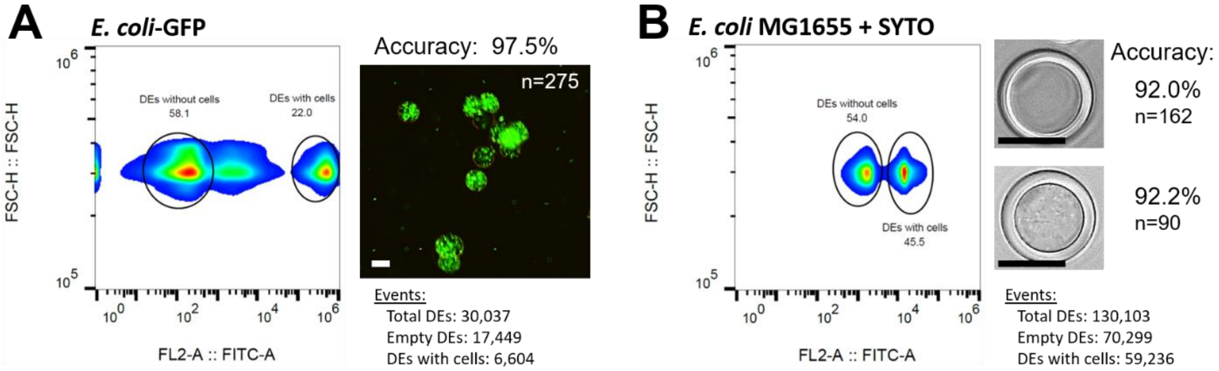


**Supplemental Figure S9. DE-FACS to sort for target bacterial populations.** FACS-sorted DEs containing grown *E. coli*-GFP (**A**) or *E. coli* MG1655 encapsulated with 2.5µM SYTO9 (**B**). Representative microscopy images from sorted 30 µm DE populations are shown, and accuracy was determined by manual fluorescence (**A**) or brightfield microscopy counts (**B**). *E. coli* was cultivated in DEs overnight (~17h) with M9 medium + 25 mM glucose at 37ºC prior to DE-FACS analysis and sorting using 130 µm nozzle size on a Sony SH800. Higher Poisson distributions were used to encapsulate *E. coli* cells in DEs to increase the populations of droplets containing cells for downstream DE-FACS and microscopy analysis. Scale bars on microscopy images indicate 20 µm.

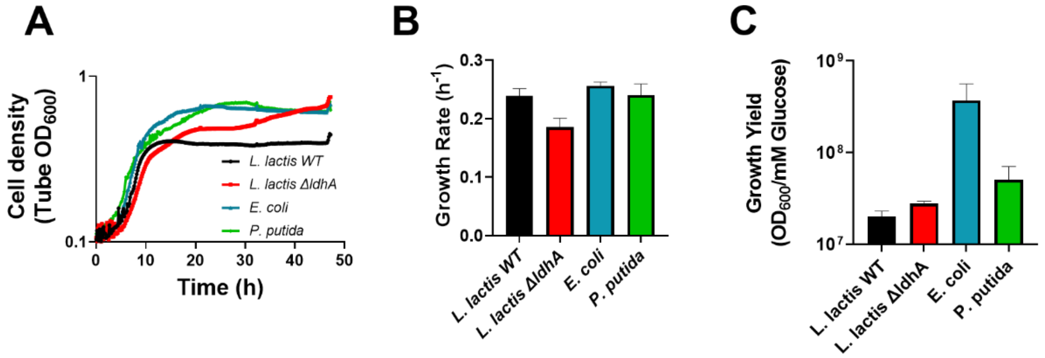


**Supplemental Figure S10. Monoculture growth trends in strains competing in mock community**. Representative growth curves (**A**), growth rates (**B**) and growth yields (**C**) of *E. coli*, *P. putida*, *L. lactis* WT, and *L. lactis* Δ*ldhA* monocultures grown in CDM + 25 mM glucose at 30°C. Error bars indicate SEM, n=3.

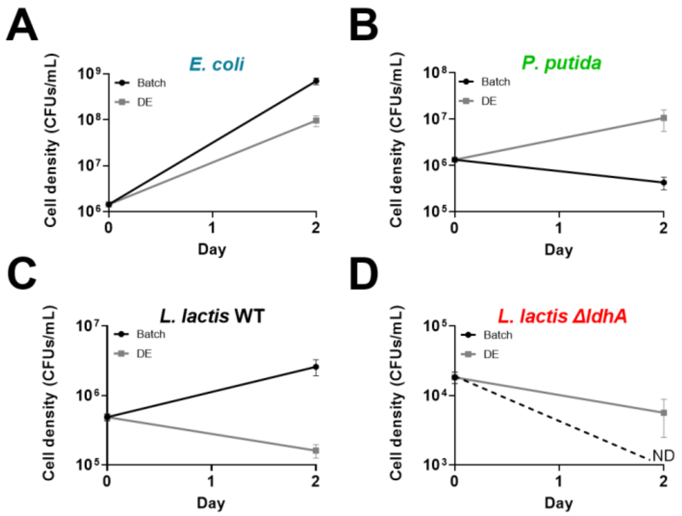


**Supplemental Figure S11. Enrichment outcomes of mock community in batch or GrowMiDE cultures.** Cell densities of mock community strains before and after enrichment in either batch or GrowMiDE cultures. All enrichments were grown in CDM + 25 mM glucose at 30ºC for 48 hours. Cell densities were determined by plating on selective and differential media. Error bars indicate SEM, n=3. ND = not detected

**Table S1: Parameters used in mathematical modeling of R and Y specialists**

| Parameter | Input values | Rate specialist R (Input values) | Yield specialist Y (Input values) |
|---|---|---|---|
| $\mu_{Max}$ | --- | 0.27 $h^{-1}$ | 0.17 $h^{-1}$ |
| $K_g$ | 0.02 mM | | |
| R | --- | $1*10^{6}$ cell $mL^{-1}$ | --- |
| Y | --- | --- | $1*10^{6}$ cell $mL^{-1}$ |
| Glu | 25 mM | | |
| Lac | 0 mM | | |
| Ace | 0 mM | | |
| EtOH | 0 mM | | |
| For | 0 mM | | |
| $Y_r$ | --- | $6*10^{6}$ cell $mL^{-1}$ mM $glucose^{-1}$ | --- |
| $Y_y$ | --- | --- | $2*10^{7}$ cell $mL^{-1}$ mM $glucose^{-1}$ |
| $F_l$ | --- | $2.25*10^{-7}$ µmol $cell^{-1}$ | 0 µmol $cell^{-1}$ |
| $F_a$ | --- | $7*10^{-8}$ µmol $cell^{-1}$ | $2.5*10^{-8}$ µmol $cell^{-1}$ |
| $F_e$ | --- | $1*10^{-9}$ µmol $cell^{-1}$ | $2.5*10^{-8}$ µmol $cell^{-1}$ |
| $F_f$ | --- | $7*10^{-8}$ µmol $cell^{-1}$ | $4*10^{-8}$ µmol $cell^{-1}$ |

**Table S2: Stool Enrichments in GrowMiDE conditions**

| Condition Name | Explanation | # Samples |
|---|---|---|
| mBHI A and B | Standard enrichments in mBHI with freshly-collected stool (biological replicates) | 6 |
| mBHIplus | mBHI modified to include sugars, short chain fatty acids, and sodium bicarbonate | 3 |
| Frozen Stool | Cells were extracted from frozen stool samples | 3 |
| PBS | Cells were suspended in PBS in syringes during DE generation to prevent growth during long droplet collection times | 1 |
| MineralOil_PBS | Droplets were generated using conditions outlined in PBS, and mineral oil was overlaid on the bulk droplet pellet | 3 |

**Supplementary Information References:**
